# Dynamic exchange of antimicrobial peptides stabilize persistent lipid pores

**DOI:** 10.1101/2025.09.18.677094

**Authors:** Jason Sengel, Bing Zan, Marc Szabo, Jakob Ulmschneider, Mark I. Wallace

## Abstract

Antimicrobial peptides (AMPs) are a promising broad-spectrum complement to conventional antibiotics. But despite their relatively simple structures, the reported modes of action of AMPs are diverse, ranging from formation of stable ion-channel-like structures, or disruption of biochemical pathways, to comprehensive membrane solubilization. This apparent complexity of function is a significant hurdle in their future application and rational design. Here, we combine single-molecule tracking with optical single-channel recording in synthetic mimics of bacterial membranes to image the pores formed by AMPs in real time, mapping the diffusion of individual AMPs relative to sites of membrane permeablization. Using alamethicin, magainin-II and melittin as archetypes of three principal membrane-disrupting AMP classes, we observe AMP molecules freely diffusing on the membrane, but with an enhancement of local peptide surface density within a nanoscopic region surrounding each pore locus. We do not observe this enhancement for indolicidin, an AMP which is not believed to use pore formation as its main mode of toxicity. Corroborated by molecular dynamics simulations that replicate our experiments, these results suggest defects formed by membrane-active AMPs are persistent but dynamic structures, stabilised by individual peptides free to diffuse into and out of the pore. Whilst alamethicin, magainin-II and melittin are often placed in different mechanistic pore-forming groups, the broad similarities between observations in these different AMPs encourages a continuum view of pore-forming ability, rather than discrete categories of mechanism.

## Introduction

Antimicrobial peptides are short (<50 amino acids) amphipathic biomolecules produced by every class of life that show selective activity against bacteria, viruses, and fungi [1–5]. AMPs form part of an organism’s innate immune response [6], but are also employed in the defensive skin secretions of amphibians and the venoms of bees and scorpions [1, 6]. Their anti-bacterial activity has gained particular attention in light of the increasing problem of antimicrobial resistance to antibiotics; the ability of AMPs to selectively target a pathogen without reliance on a specific target biomolecule, along with their apparent range of mechanisms, make them an attractive potential alternative to conventional antibiotics and other therapeutics [2, 5, 7]. However, the full mechanism of action of many AMPs remains unclear, warranting further study.

AMPs exhibit broad-spectrum activity through a variety of mechanisms [3]: some attack membranes of target cells directly, forming transmembrane pores that result in loss of cytosolic material, or by mediating depolarisation or lipid clustering [3, 4]; others translocate the membrane and subsequently interfere with intracellular machinery [3, 5]. Here, we focus on AMPs whose modes of action are thought to be based upon the formation of transmembrane pores. There are three major established classes of activity for pore-forming AMPs (Fig. 1) [1–5, 8]: (a) the barrel-stave model, in which peptides line the edge of a cylindrical pore, with the hydrophobic lipid tails shielded from the aqueous lumen; (b) the toroidal pore model, in which AMPs stabilise a defect in which the lipids re-orientate to form a high-curvature torus in which lipid head-groups and peptide line the pore; (c) the carpet model, where AMPs encourage direct loss of lipid molecules from the membrane, effectively solubilising the bilayer.

**Fig. 1.**
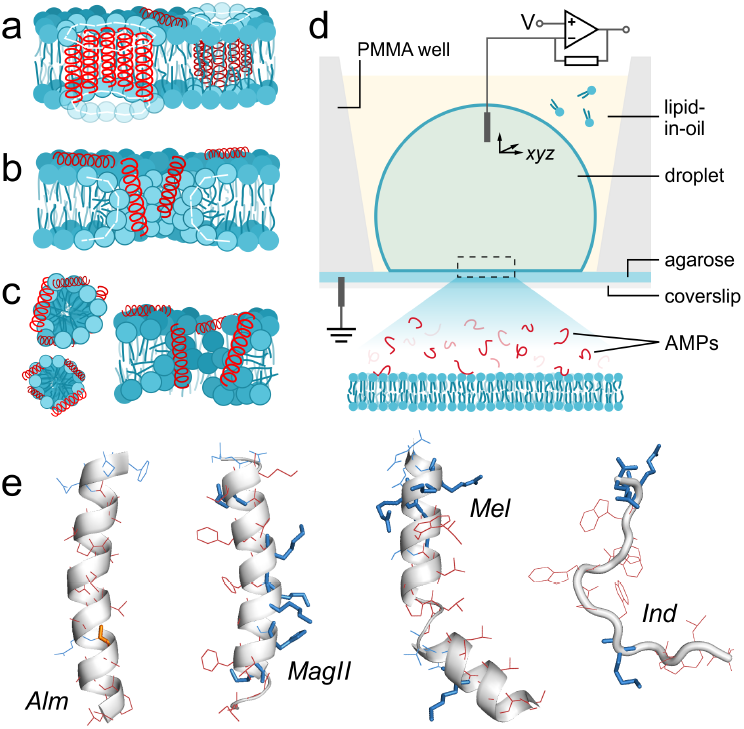
Pore-forming modes of action of AMPs. (a) Barrel-stave, (b) toroidal pore, and (c) carpet models. Lipids are in blue and AMPs drawn as red helices. Dashed white lines highlight the geometry of the different pore boundaries. (d) Schematic of droplet-interface bilayer device. A bilayer is formed between an aqueous droplet and an agarose substrate in a lipid-in-oil solution contained in a poly(methyl methacrylate) (PMMA) microfabricated chamber. AMPs present in the droplet insert into the bilayer from above. Transmembrane voltages are applied through a patch-clamp amplifier, with recording facilitated by electrodes in the agarose and in the droplet. (e) AMPs used in this work (PDB structures 1AMT, 2MAG, 2MLT and 1G89). The C-terminus is shown uppermost. Colour coding: *blue*, hydrophilic side chains; *red*, hydrophobic residues; *orange*, residues that differ in *Alm* homologues. Thicker side chains show those that bear a charge.

Although pore-forming AMPs have been subject to extensive study, much of our current mechanistic understanding relies on linking static structural experiments to ensemble measurements of function. The weakness of these approaches is that they do not tackle the core behaviour at the molecular level - namely, can we understand directly how a localised membrane defect is formed by an AMP?

Single-molecule measurements of membrane disruption have the potential to make the link between molecular action and a fluorescent readout of membrane disruption. Fluorescence microscopy has been highlighted as a powerful tool for the study of antimicrobial peptides, but studies generally measure the response of whole cells or assay ensemble vesicle leakage [9]. Pioneering single-molecule AMP experiments include the use of single-molecule Förster resonance energy transfer to track the interaction between oncocin and actively translating ribosomes [10], tracking AMPs on the surface of supported lipid bilayers [11], and the use of surface-induced fluorescence attenuation to infer the membrane orientation of LL-37 [12, 13]. However, to our knowledge a direct experimental link between the dynamics of pore formation and the action of AMP peptides has not been made. To address this, we applied singlemolecule fluorescence microscopy to image both AMP-mediated transmembrane defects and the peptides that formed them. Using droplet-interface bilayers (DIBs, Fig. 1d) [14, 15] with lipid compositions designed to mimic bacterial membranes, we tracked fluorescently-labelled AMPs in real time, correlating their location on the membrane with the location of individual transmembrane pores, which were visualised using a fluorescence reporter of calcium flux through the pore [16]. We chose to study three AMPs that broadly represent each of the three pore-forming categories: alamethicin (*Alm*, barrel-stave), magainin-II (*MagII*, toroidal pore) and melittin (*Mel*, carpet model). As a potential control, we also selected indolicidin (*Ind*) which is not thought to permeabilise membranes directly. Secondary structures are shown in Fig. 1e, and to assist readers unfamiliar with these AMPs, we first provide a brief overview of current knowledge:

*Alm* is a 20-residue peptide extracted from the fungus *Trichoderma viride* that demonstrates broad-spectrum antibacterial activity at micromolar concentrations [17, 18]. It forms a kinked *ω*/3_10_-helical structure [19], associating parallel to membrane surfaces and thinning the membrane at low concentrations [20, 21] but reorientating perpendicular to the bilayer plane at higher peptide-to-lipid (P/L) ratios [22–24]. Forming discrete conductance states in membranes [25, 26], the most widely-accepted mechanism of alamethicin’s toxicity is the barrel-stave model (Fig. 1a), in which multiple *Alm* molecules insert perpendicular to the membrane surface to line a water-filled pore [19, 27], with the discrete conductance steps observed due to changing numbers of monomers lining the pore.

*MagII*, found in the skin secretions of frogs of the *Xenopus* genus, is a 23-residue, cationic (+4) peptide that demonstrates antibacterial activity against a range of Gram-positive and -negative species at micromolar concentrations [28, 29]. *MagII* is unstructured in solution [30, 31], but adopts an *ω*-helical structure when associated with bilayers, orientating parallel to the bilayer surface [31–33] and thinning the membrane [21, 34]. Like *Alm*, increasing the P/L ratio results in a greater proportion of *MagII* adopting a transmembrane orientation [34, 35]. *MagII* can dissipate transmembrane potentials [36], induce leakage of molecules from anionic liposomes [32], and cause channel-like disruption, with a large variability in the amplitude of the current fluctuations [37]. Permeabilisation is thought to be *via* toroidal pores (Fig. 1b), where both lipids and peptide line the pore. This is evidenced by neutron scattering data indicating that only 4 to 7 *MagII* monomers are present in pores, larger than those formed by *Alm* [38], and that *MagII* accelerates the half-life of lipid flip-flop from hours to minutes in asymmetric vesicles [39]. These observations are consistent with a toroidal pore geometry presenting a route for rapid inter-leaflet lipid exchange *via* the curved pore walls.

*Mel*, a major component of honey bee (*Apis mellifera*) venom, inhibits bacterial and fungal growth, displaying micromolar minimum inhibitory concentrations (MICs) and strong haemolytic ability [40–43]. Despite having a similar structure and charge (+6) to *MagII, Mel* shows a lytic activity towards zwitterionic bilayers two orders of magnitude stronger than *MagII* [44]. Unstructured in dilute aqueous solution [45], the 26-residue peptide forms a bent *ω*-helical rod [46–48], reorienting to a transmembrane state as the P/L is raised [49]; once inserted, *Mel* causes leakage from vesicles [50] and induces increasing bilayer current fluctuations with increasing peptide concentration [51, 52]. Neutron and X-ray scattering indicates pores of comparable size to magainin pores in aligned bilayers. *Mel* is therefore reported to disrupt membranes *via* both toroidal pore model [49, 53, 54] and carpet-like manner (Fig. 1c) causing membrane fusion [55] or loss of entire bilayer patches [56, 57].

*Ind* is a short cationic (+4) peptide produced by bovine neutrophils. Only 13 residues in length, it contains the highest proportion of tryptophan of any known peptide, with three-quarters of its residues being hydrophobic [58]. *Ind* is active at concentrations of tens of micromolar against bacteria and fungi, whilst also displaying haemolytic activity [58–60]. In buffer and in the presence of vesicles, *Ind* is disordered, and does not penetrate deeply into the lipid core [61, 62]. *Ind* has been shown to release negatively-charged dyes from PC vesicles, but not smaller, neutral molecules such as glucose [63]. Similarly, 30 µM *Ind* was only able to release 50% of an anionic dye from PC vesicles after more than 30 hours, whereas pore-forming *Mel* at 1.7 µM released all the dye within one-tenth of that time [61]. Whereas *Ind* can disrupt some model membranes [60], *Ind* was found not to lyse *E. coli* membranes at up to 16 µM, but instead to induce filamentation, a result of the inhibition of DNA synthesis [64]. Higher concentrations also slow protein synthesis. The evidences of organic ion transport across membranes, DNA binding [65] and dysregulation of internal celluar processes in bacteria suggest that *Ind* has a mainly non-lytic mode of toxicity, entering the cell *via* disruption in lipid packing [66–68].

## Results

### Electrical behaviour of antimicrobial peptides in DIBs

We first used single-channel electrophysiology to characterise the disruption of DIBs by our four representative AMPs. Peptides were used at micromolar concentrations, consistent with the local concentrations of AMP experienced by an invading pathogen, and those used in previous AMP studies [50, 69–71]. DIBs were formed from a ternary mixture of 1-palmitoyl-2-oleoyl-*sn*-glycero-3-phosphoethanolamine (POPE), 1-palmitoyl-2-oleoyl-*sn*-glycero-3-phosphoglycerol (POPG) and cardiolipin (CL) in a ratio of 69:29:2, chosen to mimic the outer membrane of bacteria [72], a common target for these AMPs.

Figure 2a presents typical single-channel behaviour in the presence of AMPs. *Alm*, at low concentrations, gated with discrete steps between conductance levels of differing heights (∼1.6 nS) due to individual peptide monomers joining and leaving the barrel-stave pore structure [25, 26, 73]. Interestingly, at concentrations above →1 µM this behaviour was less readily observed: the early signs of electrical activity were fluctuations with poorly-defined conductance, akin to the fluctuations of electropores, but at voltages well below those that generate electropores in the absence of peptide. As expected for toroidal pore-forming peptides, the electrical activities of *MagII* and *Mel* did not show well-defined, discrete conductances. Rarely, step-like changes in conductance were observed, but these activities were inconsistent, interspersed with random fluctuations, and displayed a wide range of conductance steps, from 0.2 to 2 nS. The traces produced by *Ind*-containing bilayers similarly lacked reproducible stepwise gating. However, the voltages required to observe these currents were generally larger than for the other peptides (>100 mV vs. <60 mV for the other peptides), and at a magnitude where electroporation is expected. Notably, *Ind* did show an additional minor conductance mode, characterised by transient current spikes lasting 10–20 ms, observed when baseline fluctuations were small. This type of signal has been noted in the case of cell-penetrating peptides [74], and has been suggested to be indicative of momentary defects as peptide translocates the lipid membrane.

**Fig. 2.**
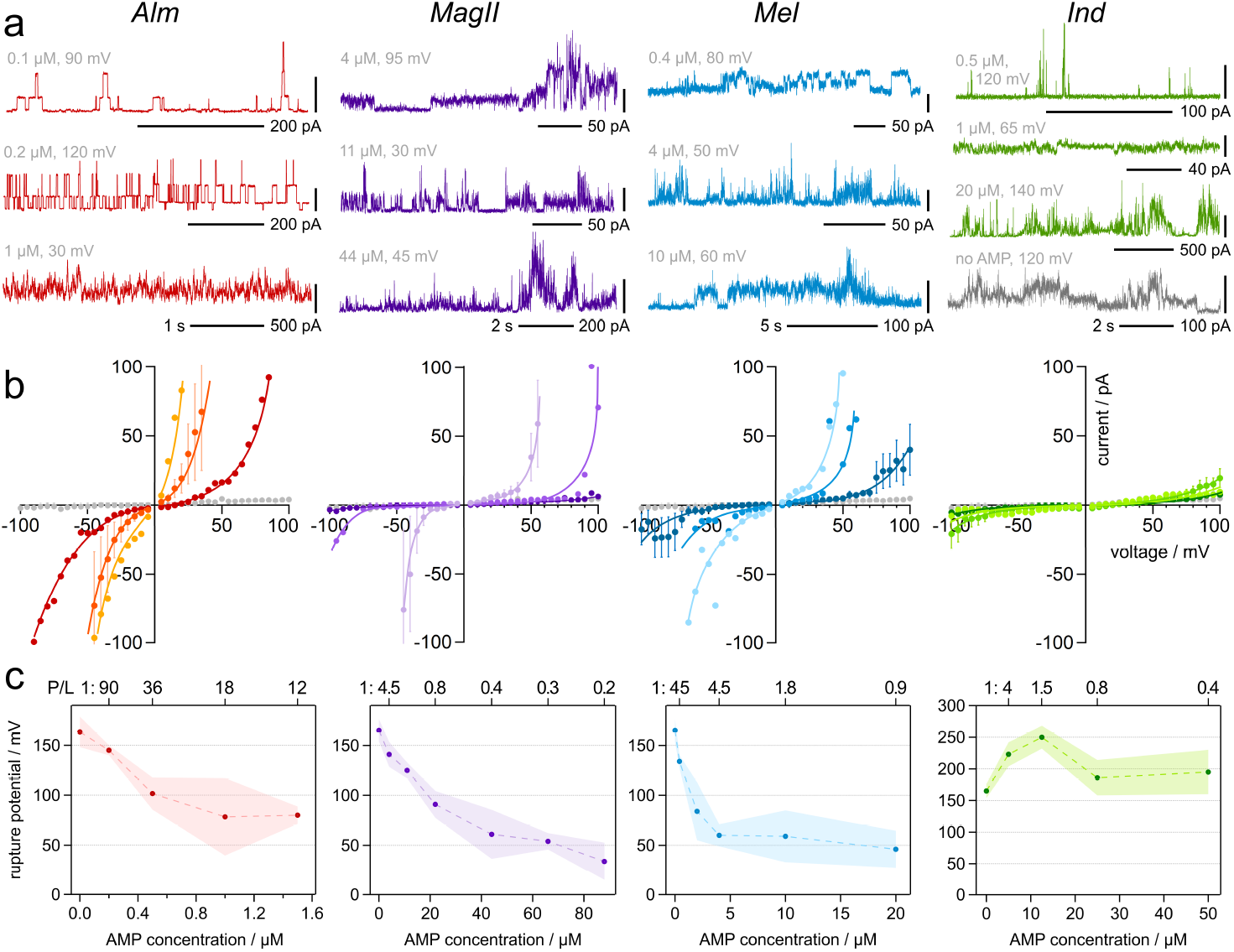
Electrical Characterisation. (a) Current recordings from AMP-containing DIBs, demonstrating a range of behaviours. *Alm* at low concentrations gates with discrete steps, but at higher concentrations of peptide discrete fluctuations are no longer resolvable. *MagII, Mel* and *Ind* show electropore-like gating, with occasional ill-defined discrete events. *Ind* exhibits rapid, transient spikes in current. A trace typical of the fluctuations occasionally recorded in plain, AMP-free bilayers (*grey*) is shown for comparison. The base of each vertical scale bar indicates the 0 pA level. (b) *I*–*V* curves (mean bilayer current vs. applied potential) for AMP-containing bilayers (coloured markers) and bilayers with no AMP (grey markers). Lighter markers indicate higher peptide concentrations, with the following concentrations displayed: *Alm* 0.2, 0.5 and 1 µM; *MagII* 4, 22 and 66 µM; *Mel* 0.4, 4 and 10 µM; *Ind* 0.5, 12.5 and 25 µM. Data points are the average of at least three independent bilayers; error bars (± 1 s.d.) are shown for one *I*–*V* curve for clarity. Lines are a guide for the eye. (c) Dependence of the rupture potential, *V*_c_, on peptide concentration, with corresponding peptide-to-lipid ratios (P/L) also indicated. Points are mean values of *V*_c_ from at least three bilayer recordings; shading shows the standard deviation of the mean.

Figure 2b shows the current responses (*I*–*V* curves) of bilayers where the average recorded current is displayed against the applied voltage. In pure lipid bilayers (i.e., in the absence of peptide, grey markers) the bilayer is essentially non-conducting. At higher voltages (>100 mV) the electric field causes the formation of electropores in the membrane, increasing the membrane conductance [75, 76]. As the number and size of pores increases, the curve becomes steeper until the bilayer ruptures [77]. Bilayers in the presence of low concentrations of AMP (darker colours in Fig. 2b) showed a similar *I*–*V* curve to peptide-free bilayers, however, the curves became non-linear at increasingly lower voltages for *Alm, MagII* and *Mel* as the concentration of the peptide was increased. These curves are of the same form as those found for these AMPs in black lipid membranes (BLMs) [37, 52, 78]. In contrast to the other peptides, the *I*–*V* curves for *Ind* showed little change in electrical activity over a wide range of peptide concentrations; indeed, increased bilayer current fluctuations (due to pore formation) on average appeared at higher voltages than in peptide-free bilayers. This is consistent with studies in which BLMs were stabilised in the presence of Ind, tolerating voltages 50 mV above other membrane-active peptides *Ind* [66].

We also looked at the relationship between peptide concentration and the voltage at which the bilayer ruptured, the critical voltage, *V*_c_ (Fig. 2c). In line with the increase in bilayer conductance with increased AMP concentration, as shown by the *I*–*V* curves, increasing peptide concentrations lead to a reduction in *V*_c_ for all but *Ind*. For the latter, a stabilising effect of the peptide on the bilayer is evident: at all concentrations the bilayer ruptured at greater voltages than in the absence of peptide.

For the other peptides, *V*_c_ levels out with increasing concentration of AMP. Taking the area of POPE-POPG lipids to be 61.5 Å^2^ [79] and a droplet volume of 350 nL, we estimate P/L ratios for each peptide concentration. We observed the greatest increases in membrane disruption (as indicated by a decrease in *V*_c_) at P/L values consistent with the thresholds at which a significant proportion of peptide transitions from membrane-parallel to membrane-perpendicular orientations, above 1:40 in the case of *Alm* [22] and around 1:15 for *Mel* [49]. It is interesting to note that *Alm* showed destabilising activities at far lower concentrations than *MagII* and *Mel*, indicating that the overall peptide charge is not necessarily the dominating factor in the interaction between the peptide and the membrane. Also of note is that in bacteria the MIC of *Mel* was found to be between 0.2 and 0.1 of *MagII*, depending on the species that the AMPs were interacting with [29]; similarly in our experiments, *Mel* induces a given *V*_c_ at concentrations 0.1 to 0.2 times lower than *MagII*.

### Single-molecule and optical single-channel imaging of AMP pores

In order to better understand the relationship between the AMPs and the pores formed in the membrane, we combined single-molecule fluorescence tracking and optical single-channel recording (oSCR, Fig. 3a) [77, 80]. The droplet solution contained an appropriate concentration of AMP so as to induce frequent and long-lived pore formation (*ca*. 500 nM for *Alm*, 25 µM for *MagII* and 10 µM for *Mel*). *Ind* was incorporated at 15 µM, which is above or at least the concentration found to release molecules from vesicles [63] or interfere with bacterial DNA synthesis [64]. In all cases, 2 nM of C-terminally tetramethylrhodamine (TAMRA)-labelled peptide was also added to track individual AMPs.

**Fig. 3.**
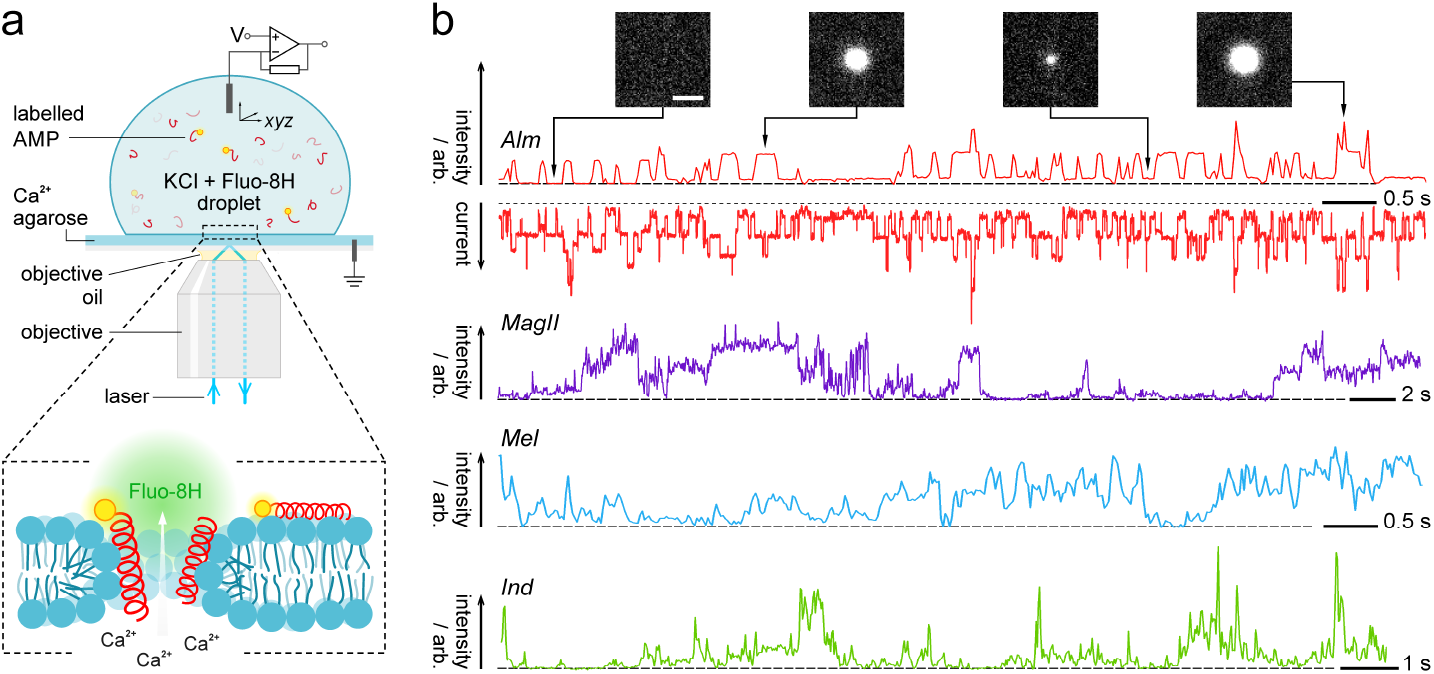
Optical single-channel recording of AMPs. (a) Total Internal Reflection Fluorescence imaging for singlemolecule and Ca^2+^-flux imaging. Fluorogenic Fluo-8H provides a localised signal corresponding to the pores in the membrane. Tetramethylrhodamine-labelled AMPs allow tracking of the location of individual molecules on the membrane. (b) Representative oSCR from AMPs, showing Ca^2+^ flux vs. time traces for peptide pores. Simultaneous electrical and optical signals are shown for alamethicin, with oSCR fluorescence (arb. units) as upward deflections and the total bilayer current as downward deflections, showing the contribution to the electrical activity by the selected pore. oSCR images are single 20 ms frames. Scale bar: 10 µm. Dashed grey lines indicate the background fluorescence intensity or 0 pA level.

To visualise the ion flux through transmembrane pores, the droplet contained the calcium-sensitive dye Fluo-8H. Upon pore formation, Ca^2+^ in the agarose is driven into the droplet and chelated by Fluo-8H, which becomes fluorescent, thus providing an optical readout of both the pore location and its individual gating behaviour under total internal reflection fluorescence (TIRF) illumination (Fig. 3b). For *Alm*, the fluorescence traces of isolated pores exhibited the same discrete fluctuations as the electrical recordings [16], showing that the step-wise changes are the result of individual pores changing in radius in a discrete manner, consistent with *Alm* monomers entering and leaving an exclusively peptide-lined pore. However, as expected, *MagII, Mel* and *Ind* showed non-stepwise fluctuations in the intensity of isolated pores.

Individual TAMRA-labelled peptides were observed as bright, mobile spots in the bilayer (Fig. S1a). Following localisation and tracking, lateral diffusion coefficients were computed from mean-squared displacement (MSD) vs. !*t* plots (Fig. S1b). In order to compare the dynamic behaviour of the peptides to that of the surrounding lipids, experiments were also carried out using 3 nM Atto590-DOPE, yielding a modal lateral diffusion coefficient (*D*_lat_) of 3.3 µm^2^ s^∼1^. The diffusivities of the four AMPs were slightly slower, as expected for membrane-associated proteins [81]; with *MagII, Mel* and *Ind* exhibiting modal values of 2–2.4 µm^2^ s^∼1^ (Fig. S1c). These values are consistent with other studies that measure the diffusivity of membrane-associated AMPs [11, 16]. *Alm* displayed a (modal) *D*_lat_ closer to that of the lipids (2.9 µm^2^ s^∼1^), which could reflect this AMP’s lower charge, with stronger association of the three cationic AMPs with the anionic membrane leading to reduced diffusivity for *MagII, Mel* and *Ind*.

Labelled AMPs diffused freely across the entire surface of the membrane; with no obvious association of individual AMP molecules with a specific pore. In order to investigate this more quantitatively, RDFs were computed relative to the location of the pore (Fig. 4). *Alm, MagII* and *Mel* all show an enhancement in the RDF, g(*r*), near the pore - within a radius of around 500 nm. RDFs computed at locations away from the pore show no enhancement (Fig. S3). The RDF for *Ind* also showed no such enhancement. As a further control, the RDF for labelled lipid around an electropore was computed (Fig. S4); with no enhancement of surface concentration around the pore locus.

**Fig. 4.**
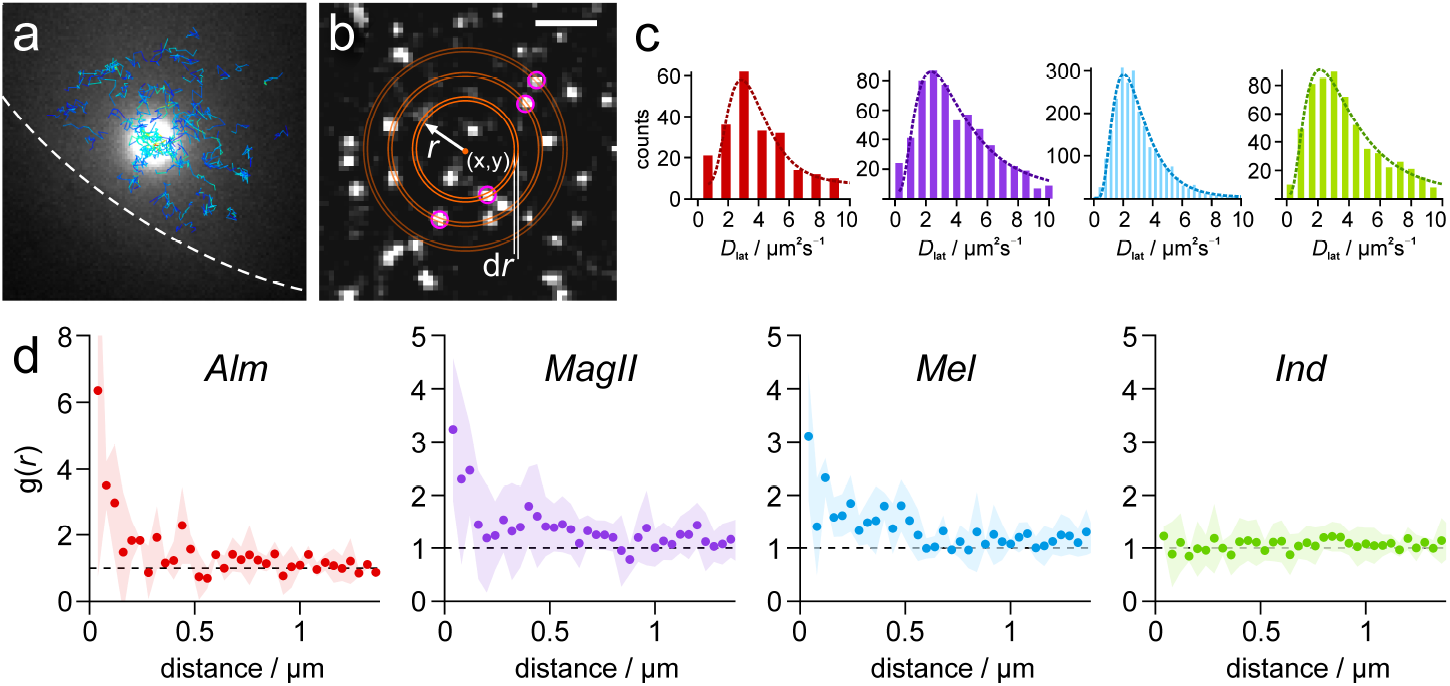
Single-molecule analysis of AMPs and their pores. (a) Blue-excitation TIRF image showing the Ca^2+^ flux through a single pore (bright white circle) in an AMP-treated bilayer. Overlaid (*blue*) are the computed trajectories of single AMPs. Dashed line is the bilayer edge. (b) Schematic outlining the RDF calculation. Single molecules within a shell of thickness *dr* are counted over the surface of the bilayer, starting at a given locus *(x, y)*. Pink circles are spots (melittin) detected by single-particle tracking. Scale bar: 5 µM. (c) Lateral diffusion coefficient (*D*_lat_) histograms for labelled AMPs. Dashed lines are log-normal fits to the data. Data was from between 3 to 5 separate bilayer recordings and between 261 (*Alm*) and 2473 (*Mel*) single molecule tracks. (d) RDFs for AMPs, computed about the pore locus. Each RDF is the mean of at least three separate pores (*Alm* = 3, *MagII* = 5, *Mel* = 3, *Ind* = 5). Shaded region is the standard deviation of the mean. Pore-forming AMPs *Alm, MagII* and *Mel* showed enhanced probability of finding peptide near the pore locus; indolicidin showed no enhancement.

### Molecular dynamics simulation of AMP location vs. pore formation

To gain further insight into our observed enhancement in AMP concentration at the pore locus we conducted molecular dynamics (MD) simulations using the methods we recently reported [82], with bilayer compositions and P:L ratios matching our experiments. Similar to our previous findings, *Alm* simulations show spontaneous insertion of peptides into the membrane, and the formation of dynamic helical bundles containing 4-8 peptides (Fig. 5a). Channels were often seen to collide, but without the exchange of peptides. Instead, the simulations revealed that bundles change size in single-peptide steps. These observations are consistent with our single-molecule fluorescent and optical single-channel recordings, where we observe discrete conductance steps from individual pores diffusing on the bilayer.

**Fig. 5.**
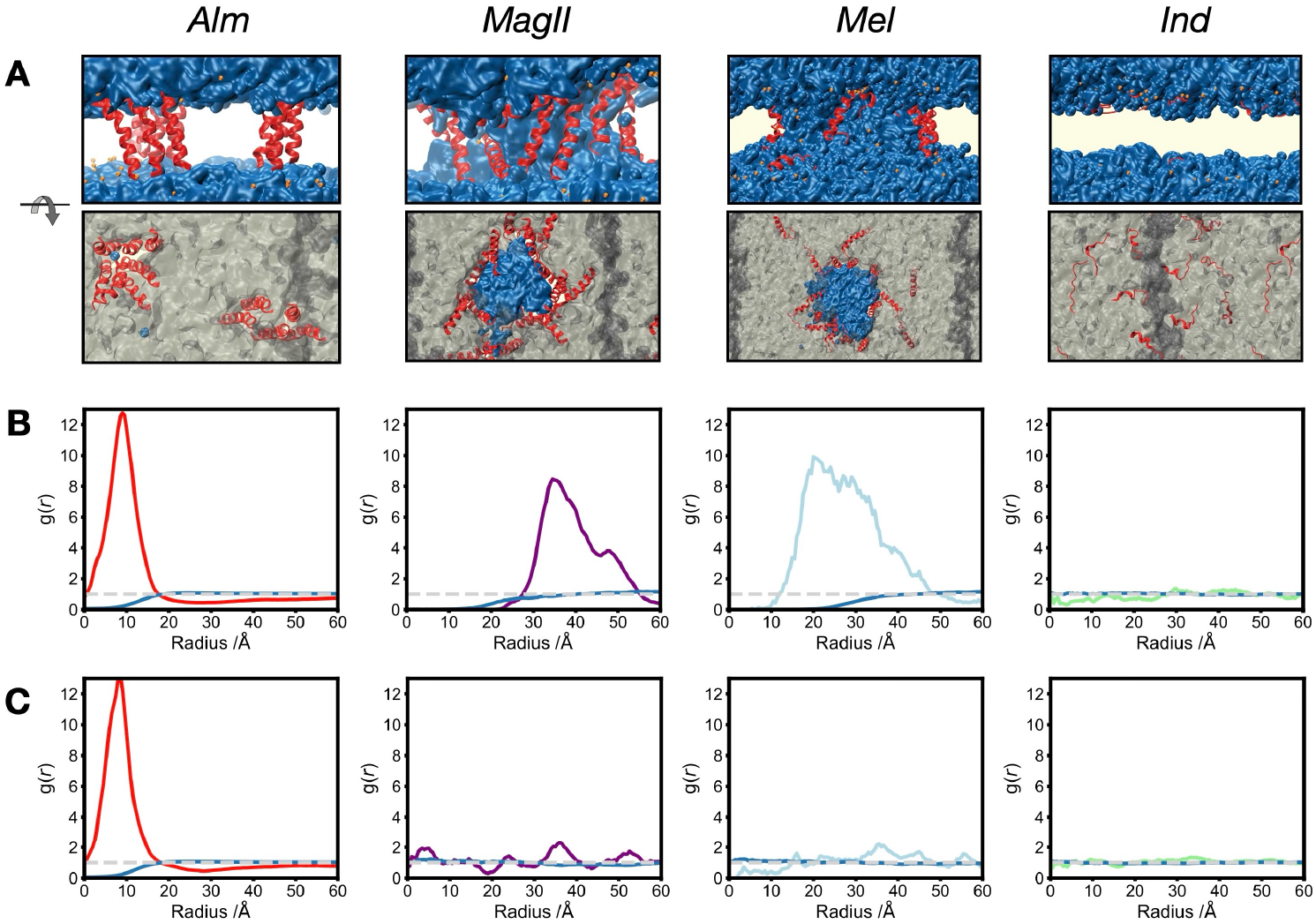
Molecular dynamics simulation of pore formation by AMPs. Simulation of all AMPs were performed in a PE:PG:CL lipid bilayer matching our experimental conditions. a) Snapshots from stable AMP pores under an applied field of 50 mV/nm 0.25 µs after simulation start. b) Radial Distribution Functions (RDFs) under an applied field of 50 mV/nm were calculated relative to the pore locus. Dark blue lines show the equivalent RDF for lipids relative to the pore locus. c) Radial Distribution Functions (RDFs) after removal of the applied field. Dark blue lines show the equivalent RDF for lipids relative to the pore locus.

In contrast to *Alm*, our other AMPs did not spontaneously insert into the membrane or form pores throughout our standard simulations (Fig. S6). We reasoned that these peptides likely permeabilise membranes on timescales much longer than achievable by equilibrium MD, and that the membrane potential present in our experiments might be required for effective pore formation by these AMPs. To address this, we used biased molecular dynamics to apply an electric field perpendicular to the membrane. Direct ab-initio prediction of assembled pores was not possible because these peptides did not spontaneously assemble from a simple transmembrane-inserted starting point. As these AMPs are thought to form toroidal pores, simulations were instead started from a pre-assembled toroidal pore (Fig. S5) in which both lipids and peptide line a substantially large water channel. A potential of 140 mV nm^∼1^ was applied for 200 ns before reducing the potential to 50 mV nm^∼1^ for the remainder of the simulation (Fig. S6). This procedure generated stable longlived electropores in the membrane (0.3 µs *MagII*, 0.8 µs *Mel*). For *Mel* and *MagII*, pores are partially lined with peptide but the majority of peptides remain on the membrane surface; in contrast to the helical TM bundles seen for *Alm*. Upon cessation of the applied potential, pores were observed to collapse (Fig. S6). For our *Ind* control, the constructed pore was observed to immediately collapse, with peptides returning to the surface of each leaflet. Simulations at 0 and 60 mV nm^∼1^ showed no pore formation, and no ion leakage was generated across the membrane. At higher potentials (140 mV nm^∼1^), a pure lipid bilayer ‘electropore’ was formed, without any *Ind* peptide involvement.

RDFs were calculated from the MD trajectories for each AMP in an analagous manner to our experiments, computing the radial distribution of peptides over the course of the simulation relative to the location of the pore (Fig. 5d). As expected from the structured pores present in the simulation, *Alm* exhibits a strong peak at small radii (*r* =1 to 2 nm) in either the presence (Fig. 5b) or absence (Fig. 5c) of an electric field. In contrast, *MagII* and *Mel* both show an enhancement in local AMP concentration relative to the location of the pore in the presence but not absence of an applied field. The radius associated with this enhancement is also larger, appearing at around 2 to 4 nm. We also computed RDFs not for the AMPs relative to the pore locus, but also of lipids (Fig. 5b&c, dark blue lines); here we see a decrease in the RDF at small radii consistent with a pore. As expected, *Ind* did not exhibit a peak in the RDF.

## Discussion & Conclusions

We have combined single-molecule fluorescence tracking with optical single-channel recording to directly observe the relationship between antimicrobial peptide location and pore formation. The central observation is that pore-forming AMPs show enhanced local surface density around active pore sites whilst retaining free diffusion across the membrane. This supports a model of dynamic pore structures stabilised by peptide exchange, rather than static assemblies with fixed stoichiometry.

Single-molecule tracking first revealed the fundamental mobility characteristics of membrane-associated AMPs. The observation of two-dimensional diffusion of fluorescently-labelled AMPs indicates that they are indeed largely surface-associated. The peptides were mobile, having diffusion coefficients slightly lower than those of labelled lipids, as might be expected for small, membrane-associated species [81]. The slightly reduced diffusivity for the cationic AMPs (*MagII, Mel*, and *Ind*) compared to *Alm* likely reflects stronger association of these charged peptides with the anionic membrane surface. These differences are perhaps inconsistent with a picture in which the majority of *Alm* peptides are transmembrane, as predicted by our simulations, as one might expect a lower diffusion co-efficient for transmembrane peptides as compared to membrane-associated molecules.

The key test of our approach was whether we could detect spatial correlation between AMP location and pore activity. RDF analysis gives clear evidence of local association of AMPs with transmembrane pores. The increased probability of finding peptide molecules about the pore locus is expected for pore-forming AMPs such as *Alm, MagII* and *Mel*. DIBs containing *Ind*, along with lipid-only bilayers, showed little discernible enhancement. The quantitative nature of the RDF analysis also allows us to compare peptide organisation at pore sites across different AMP classes. Since g(*r*) is a normalised count of the number of particles at a given distance, the relative peak values give information about the quantity of peptide in the pore. It is interesting to note that *MagII* and *Mel*, both thought to operate through the same mechanism of forming toroidal pores, both give g(*r*) ≃ 3 as the pore is approached. On the other hand, *Alm* shows a g(*r*) of approximately double this value near the pore, indicating twice the probability of finding peptide in the region relative to *MagII* and *Mel*. Assuming pores of approximately similar size between peptides (reasonable given that the oSCR signal intensity was similar across the AMPs), this is consistent with *Alm* forming barrel-stave pores in which the entire circumference of the pore is lined with peptide, and *MagII* and *Mel* forming toroidal pores, which are lined with both peptide and lipid molecules.

These quantitative differences in peptide organisation point to distinct pore architectures that nevertheless share common features. By nature of its geometry, a toroidal pore supported by AMP molecules does not need to have its full circumference covered in peptide. The lipid headgroups may form a significant portion of the defect (Fig. 1). In this sense, these AMPs are stabilising a toroidal pore, rather than defining it. Indeed,

Last and colleagues have argued that many AMPs and similar membrane-active peptides may operate within a shared energy landscape, with their binding to membranes serving to lower the barriers to toroidal pore formation [83]. We believe our data is consistent with this picture.

A critical question is whether the observed peptide-pore association reflects static binding or dynamic exchange. We looked carefully for evidence of single-molecule fluorescence tracks where AMPs might travel to the pore locus and remain there until they photobleach. We saw no evidence of stable oligomers on the millisecond timestcale of our measurement. Instead, AMPs diffuse freely. The RDFs indicate then that the AMPs are simply more likely to spend more time around the pore. This is consistent with the idea of a dynamic pore structure, one in which the AMPs are free to enter and leave, as opposed to a structure with a constant number of constituent peptides. Indeed, if the pores were not dynamic, once the pore circumference was ‘full’ of peptide, no more would leave or enter it. Since the TAMRA fluorophores bleach within several seconds, a non-dynamic pore would therefore quickly become (and remain) ‘dark’ for the majority of the recording, and we would predict no enhancement in g(*r*) around the pore over a recording lasting 40 s. As we *do* observe this enhancement, this supports our hypothesis that AMPs are able to diffuse in and out of pores in a dynamic manner.

These observations suggest that traditional mechanistic classifications may be overly rigid. The broad similarities in behaviour between *Alm, MagII* and *Mel* – despite their traditional assignment to different mechanistic classes – encourages a continuum view of pore-forming ability, rather than discrete categories of mechanism. Whilst the local organisation differs quantitatively, the fundamental process of dynamic peptide exchange appears common to pore-forming AMPs.

The MD simulations provide molecular-level insight into the experimental observations, despite the significant differences in temporal and spatial scales. Nevertheless the agreement for *Alm* is remarkable, confirming this is a classical barrel-stave former, and confirming single peptides join and leave existing pores by enlarging or lowering the pore diameter and thus conductance. At the elevated 363K used to accelerate pore formation in our simulation, these events happen on timescales of 50–300 ns and pores appear remarkably stable. At physiological temperatures, currently inaccessible by MD, the kinetics would be expected to be shifted to much longer ∼ms timescales, matching the discrete conductance steps observed experimentally. For the other three peptides, the experiments and simulations suggest that other transient and likely toroidal pore formation must be responsible that is voltage-dependent.

Several technical considerations merit discussion regarding the interpretation of our data. It is worth noting that *Alm* above ∼1 *µ*M showed less well-defined discrete conductance behaviour, instead exhibiting fluctuations more reminiscent of electropore activity but at voltages well below those generating electropores in peptide-free bilayers. This suggests that at higher concentrations, the discrete barrel-stave mechanism may give way to more complex, potentially cooperative pore formation processes.

The peaks in g(*r*) in our MD simulations are at a much smaller radius than the corresponding g(*r*) enhancement measured experimentally. For *r >* 2.5 nm, g(*r*) becomes essentially flat for all four peptides in the simulations. This length is much shorter than the ∼200 nm en-hancement seen in the experiments. These differences are largely consistent with the expected experimental resoltion, where our localisation precision in these experiments is ca. 40 nm (Fig. S2b&c).

To our knowledge, this is the first study combining dynamic single-molecule and calcium-flux imaging techniques to correlate the location of AMP molecules with respect to single transmembrane pores. These investigations were carried out on four well-studied archetypal AMPs. Although these results suggest a broad spectrum of similar activity for many membrane-active AMPs, clearly the next steps would be a comprehensive classification of AMPs, to examine if and how dynamic exchange is precisely associated with membrane activity. This work establishes single-molecule imaging in DIBs as a useful method to help understand existing AMPs and design future novel membrane-active peptides.

## Materials & Methods

### Materials

1-palmitoyl-2-oleoyl-*sn*-glycero-3-phospho-(1’-*rac*-glycerol) (POPG), 1-palmitoyl-2-oleoyl-*sn*-glycero-3-phosphoethanolamine (POPE) and cardiolipin (CL) were purchased from Avanti Polar Lipids and used without further purification. POPE and CL were stored as chloroform stocks between 10 and 25 mg*·*mL^∼1^; POPG was stored as a 25 mg*·*mL^∼1^ stock in 2:1 chloroform/methanol, in which it is more soluble than pure chloroform. Atto590-DOPE (1,2-dioleoyl-*sn* -glycero-3-phosphoethanolamine) was stored as a stock solution in chloroform; an appropriate volume was dried with the PE/PG/CL lipid mixture when single-molecule concentrations of labelled lipid were required. Ca^2+^-sensitive dye Fluo-8H (AAT Bioquest, California, USA) was stored as an aqueous solution at 1 mg*·*mL^∼1^. All stocks were stored at −20 ^*↑*^C. All other substances were purchased from Sigma-Aldrich, unless stated otherwise, and all aqueous solutions were made using 18.2 M” *·*cm Milli-Q water.

Unlabelled *Mel, MagII* and *Ind* were purchased from Bachem (Bubendorf, Switzerland). Unlabelled *Alm* (F50/5 and F50/7 isomers) was obtained from Sigma-Aldrich. The peptides were stored in powder form at −20 ^*↑*^C, and dissolved in Milli-Q water to appropriate stock concentrations when required. Labelled peptides were covalently bound to a tetramethylrhodamine dye (TAMRA; *ε*_em. max._ = 577 nm), and were synthesised by Lifetein (Somerset, New Jersey, USA) in the case of *Mel, MagII* and *Ind*. Labelled *Alm* (F50/7 isomer) was synthesised by Prof. Darren Dixon (University of Oxford, UK). The dyes were conjugated to the C-terminus of the peptides, in order to minimise perturbation of the peptide behaviour. The C-terminus in all four AMPs is more hydrophilic than the N-terminus, and is therefore more likely to remain at the lipid headgroup/solution interface when the protein inserts into the membrane [84]. Conjugation of the dye to this end of the molecule should therefore minimise steric interactions that perturb the lipid-peptide dynamics. The same rationale has been employed in other labelling studies of AMPs [85, 86]. Peptide sequences are detailed in the Supplementary Information (Figure S1).

### Droplet-Interface Bilayers

DIBs were prepared as described previously [15]. Briefly, 0.75% (wt/vol) ultralow gelling temperature agarose solution was homegenised at 90 ^*↑*^C, and 140 µL was spun onto a coverslip (Menzel-Gläzer, ThermoFisher Scientific). This is the substrate agarose. The coverslip was affixed to the underside of a small, custom-made poly(methyl methacrylate) apparatus (the “device”) featuring 16 wells, 1 mm in diameter. 1% (wt/vol) agarose solution containing 650 mM CaCl_2_ and buffered with 8.7 mM HEPES was flowed into the device, where it made contact with and hydrated the substrate agarose, but did not cover it at the bottom of the wells. The wells were filled with hexadecane containing PE/PG/CL in the ratio 69:29:2 at a total lipid concentration of 3.5 mg*·*mL^∼1^ and the device allowed to rest for 1 hour to allow for monolayer formation on the substrate. Meanwhile, aqueous droplets (300–400 nL) were incubated in the same lipid-in-oil solution, again for monolayer formation. The droplets contained 1.3 M KCl, 8.7 mM HEPES, 375 µM EDTA and 23 µM Fluo-8H, in addition to unlabelled (200 nM– 90 µM) and labelled (2 nM) AMP. After 1 hour of incubation, droplets were pipetted into the wells of the device where they sank onto the substrate, forming bilayers by the contact of the droplet- and agarose-associated monolayers.

Ag/AgCl electrodes inserted into the hydrating agarose, and the top of the droplet *via* a micromanipulator, allowed electrical access. Voltages were applied and bilayer currents recorded using an Axopatch 200B patch-clamp amplifier and headstage (Axon Instruments, Molecular Devices, California, USA), with signals recorded and filtered at 1 kHz. Devices were placed within a Faraday cage atop an inverted microscope (TiE Eclipse, Nikon, UK).

Bilayers in the presence of increasing concentrations of AMP were exposed to a positive and then negative voltage (with respect to the droplet) every 180 s each, before being incrementally increased by 5 mV in magnitude. Each application of voltage was separated by a rest period of 30 s at 0 mV. This cycle, starting at *±*5 mV, was repeated with increasing voltages until the bilayer ruptured. Such a protocol allows the comparison of the electrical behaviour of different bilayers subjected to the same electrical stimulus. Fluorescence imaging was carried out using a 60*↓*TIRF objective lens (oil-immersion, 1.45 NA; Nikon). Excitation of the sample and collection of fluorescent signals was carried out by the same lens. For excitation of Ca^2+^-bound Fluo-8H, a 473 nm continuous wave laser beam was used (power at the back focal plane of the objective was 1–4 mW; Vortran Laser Technology, Sacramento, California, USA), with fluorescence and excitation signals (*ε*_Fluo-8H em. max._ = 514 nm) separated by a dichroic mirror (XF2037; Omega Optical, Brattleboro, Vermont, USA), and the collected signal from the bilayer transmitted through an emission filter (Brightline bandpass 525/39 nm; Semrock, Rochester, New York, USA) in order to eliminate stray excitation wavelengths. TAMRA- or Atto590-labelled lipids or peptides were excited using a 532 nm continuous wave laser beam (power at the back focal plane of the objective was 4–7 mW; Vortran Laser Technology), with a 545 nm longpass dichroic mirror and 605/55 nm emission filter (Semrock). Images were recorded using an electron-multiplying charged-coupled device (EMCCD) camera (iXon+; Andor Technology, Belfast, UK). All experiments were conducted at room temperature of *ca*. 21 ^*↑*^C.

### Image processing and analysis

Electrical data was recorded using WinEDR electrophysiology software (John Dempster, Strathclyde University, UK), from which voltage stimulus protocols were also designed and applied. Image analyses were carried out in Fiji image-processing software [87]. Fluorescence images were corrected for laser profiles by applying a fast Fourier transform bandpass filter to remove structures larger than 30 pixels (11.769 µM), and/or by division by the median *z*-stack. Tracking of diffusing AMPs was carried out using the Trackmate plugin in Fiji, which detected spots using its Laplacian of Gaussian algorithm. Detected spots were then manually checked and filtered for quality. Numerical data was processed in Igor Pro (Wavemetrics, Oregon, USA) using custom-written procedures.

To compute RDFs, the location of a pore centre was determined by 2D-Gaussian fitting to its oSCR image. This was the locus about which the RDF was calculated. Next, the image stack of diffusing peptides recorded in the same membrane area was tracked to generate AMP location coordinates for each frame. Using this information, the RDF procedure then counted the number of particles within a shell of *r* + d*r* at a distance *r* away from the pore locus. The value of *r* was incremented in steps of 40 nm, equal to the average localisation error of 2D-Gaussian fitting to single molecules (Figure S8). The shell thickness d*r* was also set at 40 nm. The number of spots found within the shell was divided by 2*πr*d*r* + *π*d*r*^2^, to normalise for the area of the shell, since in a larger shell more particles are likely to be found. Finally, to ensure g(*r*) tended to unity (*i*.*e*. the same probability of finding a particle at any given *r*, indicating no enhanced likelihood of finding a particle), the count was normalised by dividing it by the number density, the average number of spots per frame.

### Molecular dynamics simulations and analysis

All-atom MD simulations were performed and analyzed using GROMACS 2020.4 [88]. MD simulations were using the CHARMM36 force field [89], in conjunction with the TIP3P water model [90]. Electrostatic interactions were computed using PME, and a cut-off of 10 Åwas used for van der Waals interactions. Bonds involving hydrogen atoms were constrained using LINCS [91]. The integration time-step was 2 fs and neighbour lists were updated every 5 steps. All simulations were performed in the NPT ensemble, without any restraints or biasing potentials. Water, ions, lipids and the protein were each coupled separately to a heat bath with a time constant *τ*_T_ = 0.5 ps ps using velocity rescale temperature coupling. The atmospheric pressure of 1 bar was maintained using weak semi-isotropic pressure coupling with compressibility *κ*_*z*_ = *κ*_*xy*_ = 4.6 ∼ 10^∼5^ bar^∼1^ and time constant *τ*_P_ = 1 ps. Initial systems were build as peptide/membrane/water boxes, with a salt concentration of 100 mM NaCl using CHARMM-GUI [92]. Simulations were started with transmembrane inserted peptides. For Mel, MagII and Ind, initial toroidal pores were constructed. For Alm, peptides were allowed to aggregate from initially transmembrane inserted peptides. Bilayers were designed to be symmetric, with equal number of lipids in each leaflet. Individual simulations were run for 2-4 µs.

## Authors’ Contributions

MS conducted electrical experiments; JTS performed the imaging experiments analysed the data; BZ and MU conducted all simulations; JTS and MIW wrote the paper.

## Competing Interests

The authors declare no competing interests.

## Funding

MIW is supported by the European Research Council (ERC-2012-StG-106913, CoSMiC) and the Wellcome Trust (224327/Z/21/Z). JTS is supported by the European Research Council and the Engineering and Physical Sciences Research Council.

## Notes

### Competing Interest Statement

The authors have declared no competing interest.

