## Supplementary Information for "Dynamic exchange of antimicrobial peptides stabilize persistent lipid pores"

| | |
|---|---|
| *Alamethicin* | F50/5<br>Ac– UPUAUAQUVUGLUPVUUQQPhl –COO$^-$<br>F50/7<br>Ac– UPUAUUQUVUGLUPVUUQQPhl –COO$^-$ |
| *Magainin-II* | $^+$NH$_3$– GIGKFLHSAKKFGKAFVGEIMNS –COO$^-$ |
| *Melittin* | $^+$NH$_3$– GIGAVLKVLTTGLPALISWIKRKRQQ –CONH$_2$ |
| *Indolicidin* | $^+$NH$_3$– ILPWKWPWWPWRR –CONH$_2$ |

**Table 1. Peptide sequences of the AMPs used in this work.** Colour coding: *blue*, hydrophilic side chains; *red*, hydrophobic residues; *orange*, the differing residues in the alamethicin homologues. *U* = Aib, $\alpha$-aminoisobutyric acid ($\alpha$-methylalanine); *Phl* = phenylalaninol.

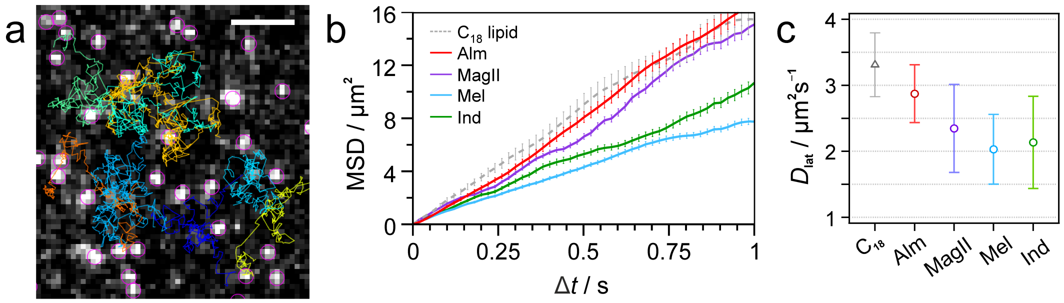


**Fig. S1. Single-molecule tracking of AMP diffusion.** (a) Green-excitation image showing single AMP molecules (melittin). Pink circles are spots detected by single-particle tracking; coloured lines show trajectories of some of the peptides. Scale bar: 5 µm. (b) Example MSD vs. $\Delta t$ plots for the four labelled peptides, along with those for the dye-labelled lipid molecule Atto590-DOPE ($C_{18}$). Error bars show the standard error. (c) Modal lateral diffusion coefficients for lipids (DOPE, *grey*) and AMPs (*colours*) in DIBs, as determined from log-normal fits to histograms of the $D_{\mathrm{lat}}$ values; error bars are the variance of the fits.

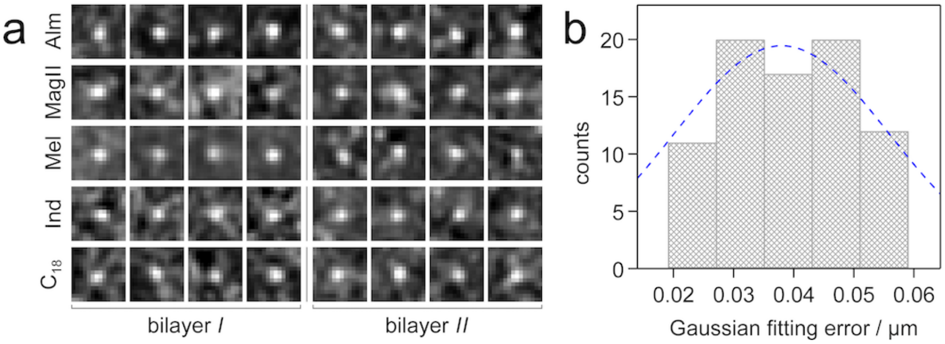


**Fig. S2.** Localisation error of 2D-Gaussian fitting to single molecules. (a) Single 20 ms frames of labelled single molecules. Four examples were taken from two separate experiments (*bilayer I* and *bilayer II*) for each petide (or labelled lipid). A Gaussian filter ($\sigma$ = 1) was applied to the images before 2D-Gaussian fitting in order to improve the accuracy of the fits. Each image is 5.49 $\times$ 5.49 µm$^2$. (b) Histogram of the errors (standard deviation) in the $x$- and $y$- loci of the 40 single molecules shown in (a), as determined by 2D-Gaussian fitting to the images. The dashed line is a fit to the data.

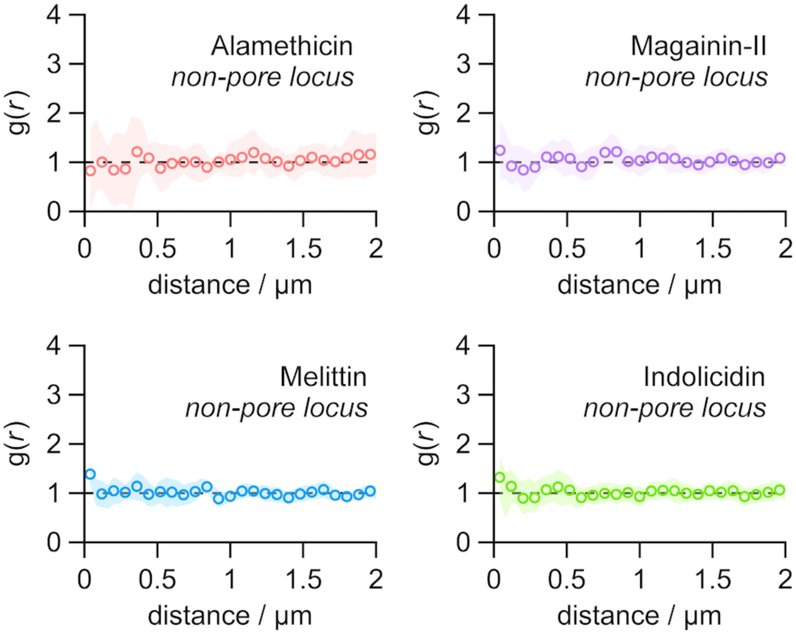


**Fig. S3. RDFs at non-pore loci for the four AMPs.** Data points are the average of at least 8 locations from multiple bilayers ( Alm = 8, *MagII* = 17, *Mel* = 12, *Ind* = 16). g($r$) remained at unity, indicating no enhancement of peptides around the reference point. Shaded region is the standard deviation of the mean.

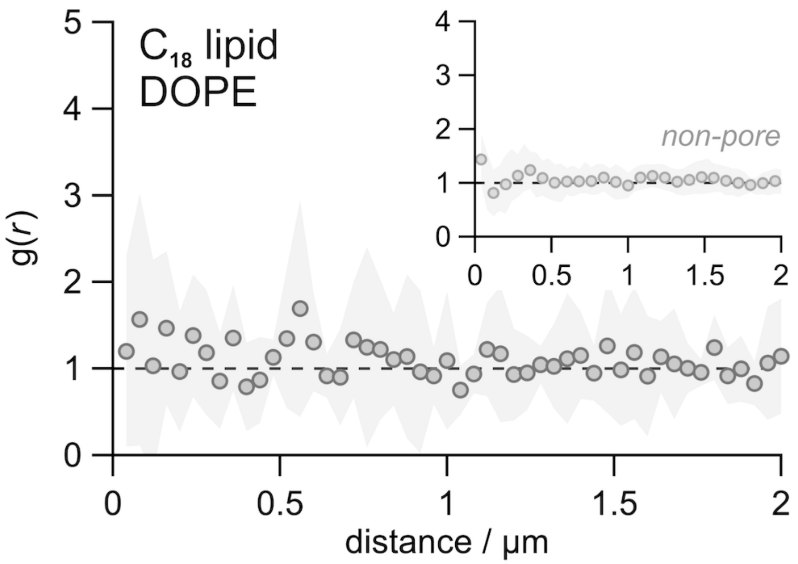


**Fig. S4. RDF controls.** RDFs for pore ($n$ = 5) and non-pore loci ($n$ = 18) for Atto590-labelled DOPE. Shaded region is the standard deviation of the mean.

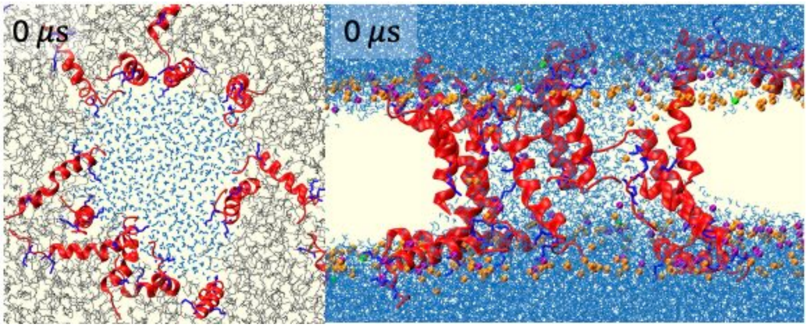


**Fig. S5. Simulation Initiation.** Simulation starting conditions for construction of Mel pores.

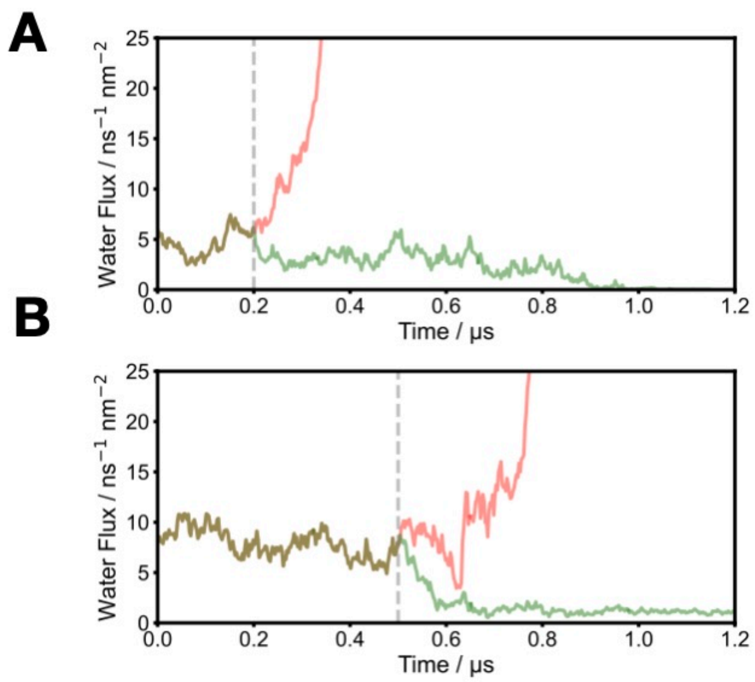


**Fig. S6. Voltage protocol for simulations** Following the 200 ns initiation procedure described in the methods, pores were relatively stable for an extended period with an external field of 50 mV nm$^{-1}$ for both (A) *Mel* and (B) *MagII*. Grey dotted lines indicate simulation bifurcation. Red indicates increasing water flux and pore expansion at 50 mV nm$^{-1}$, green indicates removal of the applied potential and subsequent pore collapse.
